# dgram2dmap: Extraction, visualisation and formatting of distance constraints from AlphaFold distograms

**DOI:** 10.1101/2022.12.08.519560

**Authors:** Björn Wallner, Alexey Amunts, Andreas Naschberger, Björn Nystedt, Claudio Mirabello

## Abstract

Distograms are data structures output by AlphaFold, along with the predicted 3D coordinates of the target protein, that encode predictions about Euclidean distances between pairs of amino acids. Although distograms are often overlooked, they do provide information that is to some extent complementary to that of the final 3D model. Here, we introduce dgram2dmap, a simple tool to convert distograms into distance maps that can be visualised and used in external tools for further downstream analyses. dgram2dmap runs within seconds to minutes, is open source and available on GitHub: https://github.com/clami66/dgram2dmap.

## 1 Introduction

Since its release, the second version of AlphaFold has revolutionised the field of structural biology by providing predictions of protein structures of unprecedented accuracy, even when no homologous structures are available in the Protein Data Bank (Jumper *et al*., 2021; Evans *et al*., 2021).

However, an often overlooked feature of AlphaFold is that, along with the 3D coordinates for each predicted model, a distogram is also provided. A distogram is a 3D matrix encoding the Euclidean distance distribution predictions for all pairs of amino acids (Senior *et al*., 2020). The distogram should not be confused with the Predicted Aligned Error (PAE), which is related to the error in pairwise distances (see Figure 1 for a comparison).

**Figure 1:**
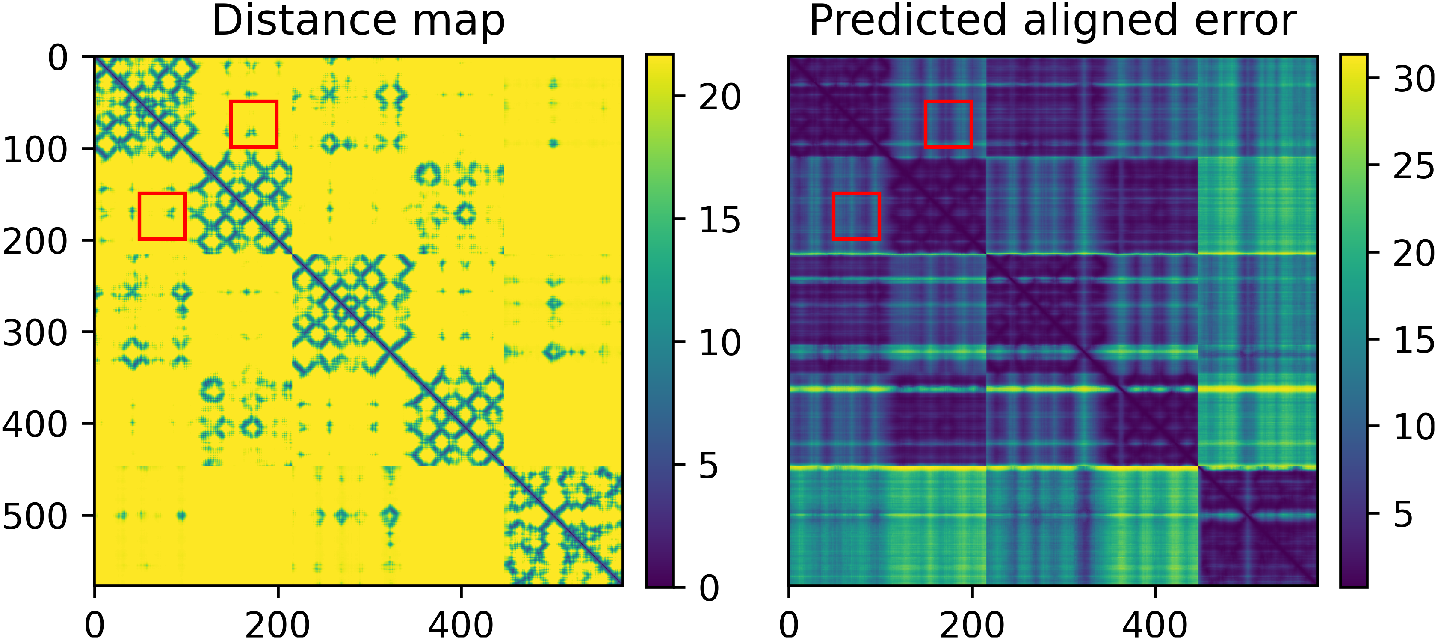
Example of dgram2dmap output plots for an AlphaFold model on CASP15 heteromeric target H1166*, where the distances between residues 50 – 100 and 150 – 200 have been highlighted (red boxes). Left: distances (Å) extracted from distogram. Right: Predicted Aligned Error plot (Å). *http://bioinfo.ifm.liu.se/casp15/H1166

Distograms cannot be easily visualised or interpreted, but are a clearer representation of the evolutionary signal that is leveraged by AlphaFold to predict a structure, and contain valuable information that is not always captured in the 3D predictions. An example of this is whenever a weak evolutionary coupling signals between distant pairs of amino acids is present, in particular across protein chains in multimer predictions (manuscript in prep.). Visualising a distogram can also be useful to analyse the nature of folding constraints for different alternative models, or investigate the evolutionary source of such constraints by using different sets of input Multiple Sequence Alignments (manuscript in prep.).

Here, we introduce `dgrap2dmap`, a tool that approximates a distogram into a 2D distance map which can then be readily visualised and compared against distance maps generated from PDB models. The distance maps can be exported in different formats for downstream analyses and calculations, e.g to Rosetta constraints for use in RosettaDock (Marze *et al*., 2018). `dgram2dmap` runs within seconds to minutes on any AlphaFold output directory containing result pickle files.

## 2 Methods

The distogram information output by AlphaFold is an array of logits with shape *N× N ×* 64, where *N* is the number of amino acids in the input sequence and 64 are distance bins, one for each range of distances that are taken into consideration when making a distance prediction (e.g., first bin has range [2.3125, 2.625) Å, the second [2.625, 2.9375) Å, and so on). The logits are normalised using a `softmax` function, similar to a classifier neural network, to obtain the probability that the final distance will fall in any given bin. The probabilities across bins for any given pair of amino acids will sum up to one. In order to get an approximate distance for a pair (*i, j*), *Dij*, the softmax output for that given pair is taken and used to perform a weighted sum of bin edges *Eb*:

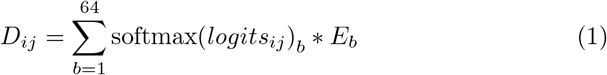

This operation might underestimate distances whenever AlphaFold is less confident about a pairwise distance, so depending on the application, it is also possible to perform an `argmax` operation instead, which will approximate the distance as the left edge of the most likely bin:

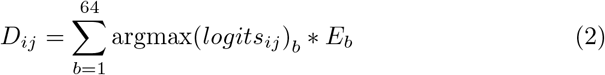

Since the highest bin limit is 21.68 Å, all distances are capped at 22 Å, even when comparing predicted distances against distances from 3D models (Figure 2).

**Figure 2:**
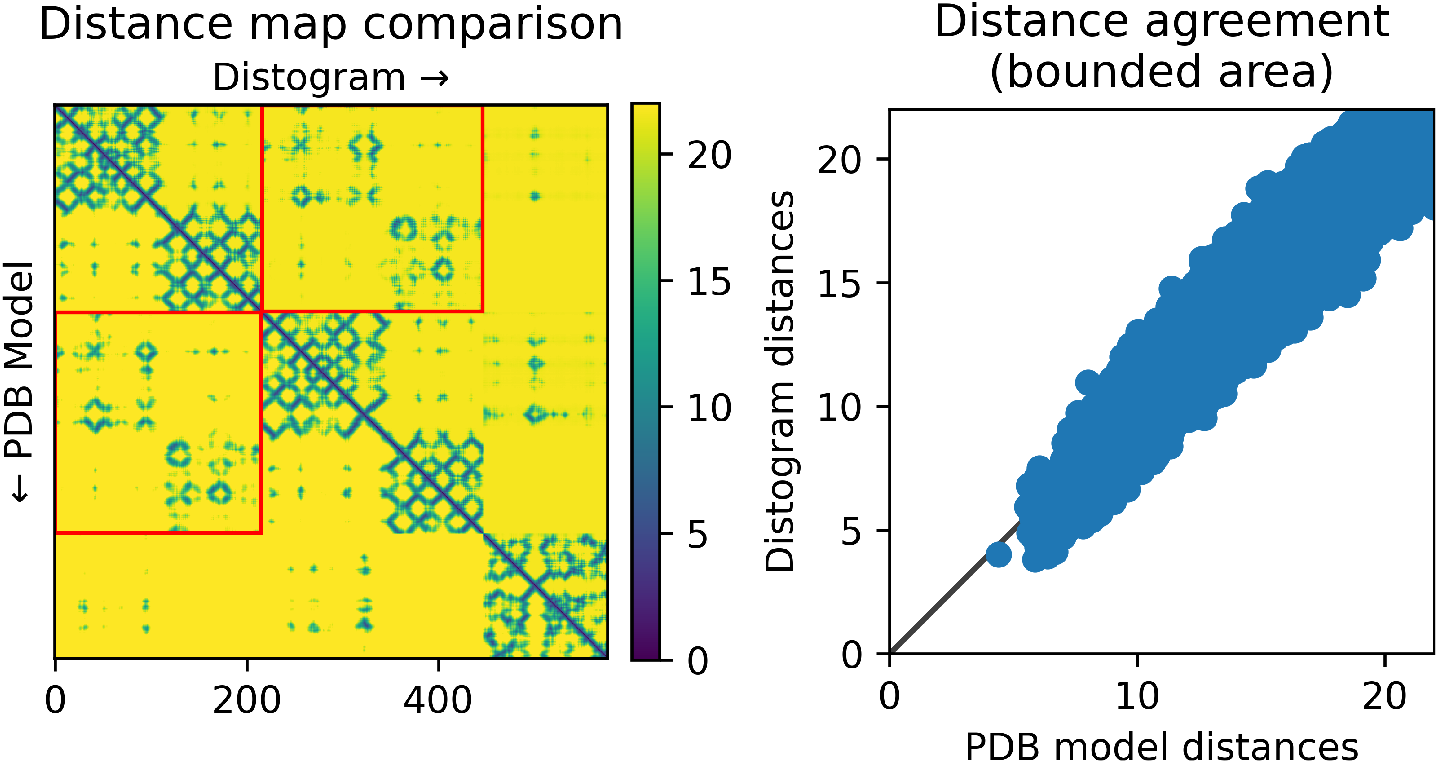
Example of dgram2dmap output plots for an AlphaFold model on CASP15 heteromeric target H1166, when compared to an AlphaFold 3D model. Left: comparison between 3D model (lower triangle) and distogram-extracted distance map (upper triangle) where the A-B interface distances have been highlighted (red boxes). Right: scatter plot of distances from red bounding boxes in distogram and 3D model. Distances are capped at 22Å.

## 3 Usage

### 3.1 Basic

dgram2dmap is a Python (Van Rossum and Drake, 2009) script that depends on *scipy* (Virtanen *et al*., 2020), *matplotlib* (Hunter, 2007), *biopython* (Cock *et al*., 2009) packages, which can be installed through *pip* or similar.

The basic way of running dgram2dmap, is through the command:

~~~
python dgram2dmap.py input_folder/
~~~

where `input_folder/` is the path to the AlphaFold output directory containing the output pickle files named `result_model_*.pkl`. The script will automatically extract and convert distograms for all models and save the distance matrices as CSV files in the input folder, with naming `result_model_*.pkl.dmap`.

### 3.2 Plotting

To visualise the distances, it is possible to run:

~~~
python dgram2dmap.py input_folder/ --plot
~~~

which will plot the distance matrices, along with the PAE, on files with naming `result_model_*.pkl.dmap.png`. If the user wants to highlight a subset of amino acids, it is possible to specify either two chains or two residue number limits as follows:

~~~
python dgram2dmap.py input_folder/ --plot --chains A B
python dgram2dmap.py input_folder/ --plot --limits 50:100 150:200
~~~

In the first case, the patch of distances corresponding to the interface between chains *A* and *B* will be highlighted with red bounding boxes (see Figure 2), in the second case, the bounding boxes will correspond to distances between two stretches of contiguous amino acids: the first between 50 and 100, the second between 150 and 200, see Figure 1. When the model is a multimer, the numbering is based on the length of the concatenated sequence of all monomers.

### 3.3 Comparing to 3D models

It is possible to perform comparisons against 3D models in PDB format, either native structures or models from any method, as long as chain names and residue numbering are consistent:

~~~
python dgram2dmap.py input_folder/ --chains A B --pdb model.pdb
~~~

which will produce PNG outputs on files with naming:

`result_model_*.pkl.agreement.png` similar to the one shown in Figure 2.

### 3.4 Exporting distances to Rosetta constraints

Subsets of distances can be extracted to Rosetta constraints files. This is useful, for example, when in multimeric predictions distograms show a possible interface between two chains that is missing in the 3D model, and the user wants to dock them in Rosetta instead:

~~~
python dgram2dmap.py input_folder/ --plot --chains A B --rosetta
~~~

This will output Rosetta constraints files with naming:

`result_model_*.pkl.rosetta_constraint`. The constraints will be between *CA* atoms with the function of type *HARMONIC*, with the mean corresponding to the predicted distance and standard deviation 1.0. The default constraint function can be configured through the `--rosetta format` flag. Constraints can also be output up to a maximum distance using the `--maxD` flag.

## Author contributions

BW developed the tool, wrote the manuscript; BN acquired the funding necessary to develop the tool, managed the project; AA and AN contributed experimental data used during the development of the tool; CM conceived and developed the tool, wrote the manuscript.

## Acknowledgements

This tool was developed within the *BeyondFold* Technology Development Project at the Science for Life Laboratory (SciLifeLab). CM is financially supported by the Knut and Alice Wallenberg (KAW) Foundation as part of the National Bioinformatics Infrastructure Sweden at SciLifeLab. BW and AA are financially supported by KAW as part of the WASP-DDLS joint program, AA is supported by the European Research Council (ERC-2018-StG-805230). Computations were performed at NSC Tetralith/BerzeLiUs provided by the Swedish National Infrastructure for Computing (SNIC), partially funded by the Swedish Research Council through grant agreement no. 2018-05973.

